# Modern botanical analogue of endangered Yak (*Bos mutus*) dung from India: Plausible linkage with living and extinct megaherbivores

**DOI:** 10.1101/388819

**Authors:** Sadhan K. Basumatary, Hukam Singh, H. Gregory McDonald, Swati Tripathi, Anil K. Pokharia

## Abstract

The study present to document the micro and macrobotanical remain on wild Yak dung to understand the diet, habitat, and ecology in relation to determining possible ecological relationships with extant and extinct megaherbivores. Grasses are the primary diet of the yak as indicated by the abundance of grass pollen and phytoliths, though it is obvious. The other associates non-arboreal and arboreal taxa namely, Cyperacaeae, Rosaceae, Chenopodiaceae, *Artemisia, Prunus*, and *Rhododendron* are also important dietary plants for their survival. The observation of plant macrobotanical remains especially the vegetative part and seed of the grasses and Cyperaceae also indicates good agreement with the palynodata. The documented micro and macrobotanical data is indicative of both Alpine meadow and steppe vegetation under cold and dry climate which exactly reflected the current vegetation composition and climate in the region. The recovery of *Botryococcus, Arcella*, and diatom was marked though in trace values and suggestive of the perennial water system in the region which incorporated through the ingestion of water. Energy dispersive spectroscopy analysis marked that the element contained in dung samples has variation in relation to the summer and winter which might be the availability of the food plants and vegetation. This generated multiproxy data serves as a strong supplementary data for modern pollen and vegetation relationship based on surface soil samples in the region. The recorded multiproxy data could be useful to interpret the coprolites of herbivorous fauna in relation to the palaeodietary and paleoecology in the region and to correlate with other mega herbivores in a global context.

## Introduction

Recently, there has been an increasing interest in the study of pollen and non-pollen palynomorphs preserved in herbivore dung and how dung can serve as a substrate for their preservation. This added information on the plant community aids in better understanding the dietary habits of herbivores in relation to local vegetation composition and climate in a region. It also provides a measure as to how dependent herbivorous animals are on the availability of different plant species within their habitat. The diversity of available plants and their relative abundance, in turn, reflects the climate of the region. The study of the modern pollen deposition on the landscape forms a critical dataset and a prerequisite to understanding the palaeovegetation and climate in the region [1–5].

The systematic study of the relationship between modern pollen and vegetation in the higher parts of the Himalayan Mountains is very difficult due to hilly terrain and limited availability of soil samples. Consequently, it may not serve as a modern analogue that would permit an accurate interpretation of the palaeoecology in the region. Previously, some workers have only examined surface soil samples in order to understand the modern pollen and vegetation relationship in the higher Himalayan region [6,7]. A complementary data set can be provided by an examination of modern herbivore dung and which can also serve as a source of modern analogues of local and regional vegetation [8–12]. It is clear that pollen and spores incorporated into the stomach contents also reflect the composition of the vegetation in relation to climate [13,14]. The distribution of herbivorous animals within an ecosystem is often dependent on vegetation composition and its regional distribution [15–18]. Study of both diatoms and phytoliths in the dung can also serve as a powerful proxy for palaeoenvironmental reconstruction and recognition of the presence of domestic herbivores [19–24].

The main aim of this study is to document the presence of both the micro and macrobotanical remains in wild Yak (*Bos mutus*) dung in order to determine the dietary preferences of the species relative to the local vegetation composition and climate. Since large animals may play an important role in the biogeochemical cycle of the ecosystem [15,25,26], we have included in this study FESEM-EDS analysis in order to determine the relationship between the elements contained in Yak dung derived from the current vegetation composition and their dietary habits in the region.

### Study sites, vegetation and fauna

The distribution of the wild Yak in India is very restricted and confined to higher elevations of the Himalaya. For this study we have selected a region which is around 12 km north from the Dronagiri village areas (Fig 1. a. & b.) in the Chamoli district of Uttrakhand (India) based on the availability of the wild Yak and accessible terrain (Fig 2).

**Fig 2.**
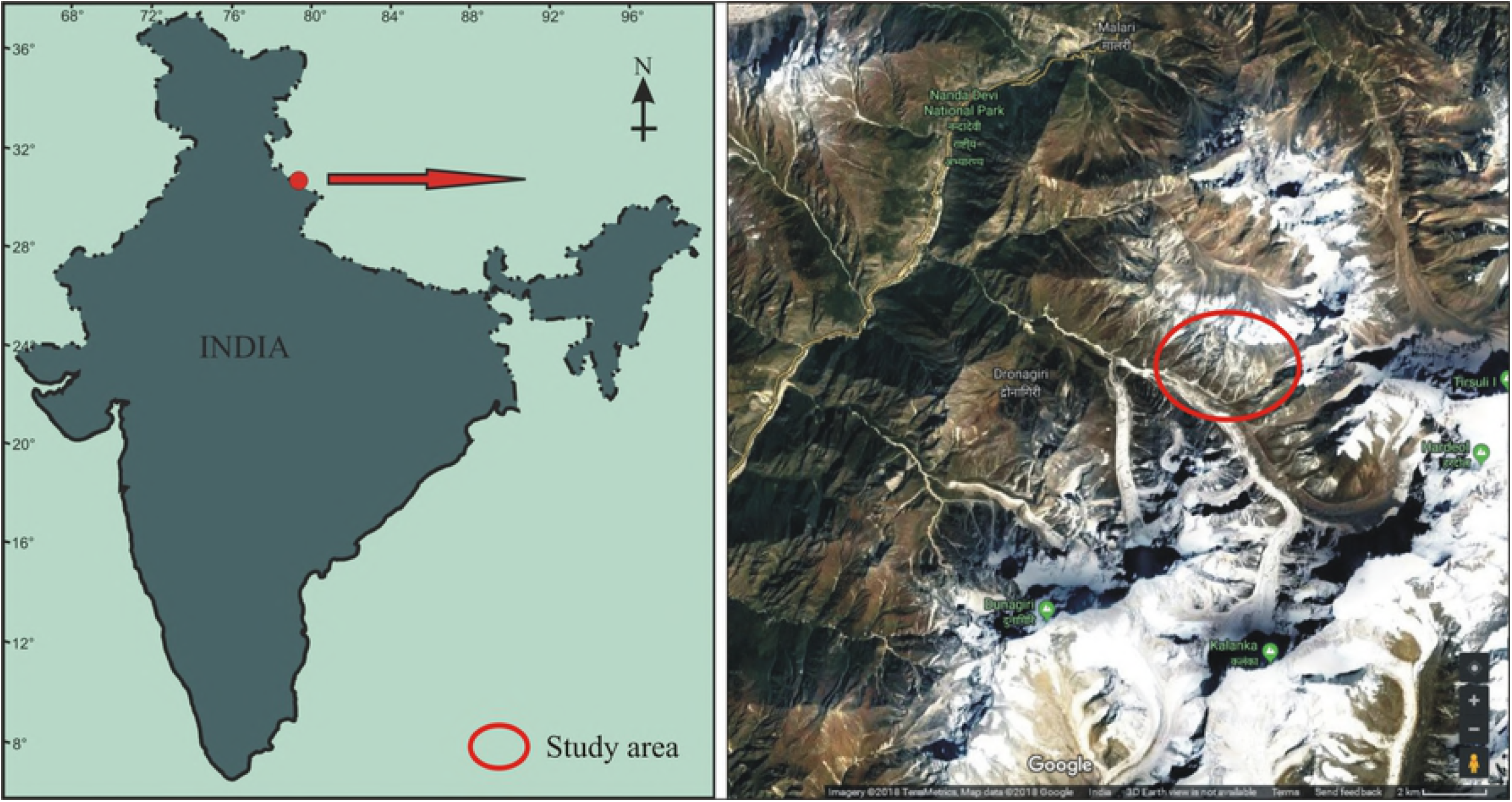
Location map of the study areas

**Fig 1.**
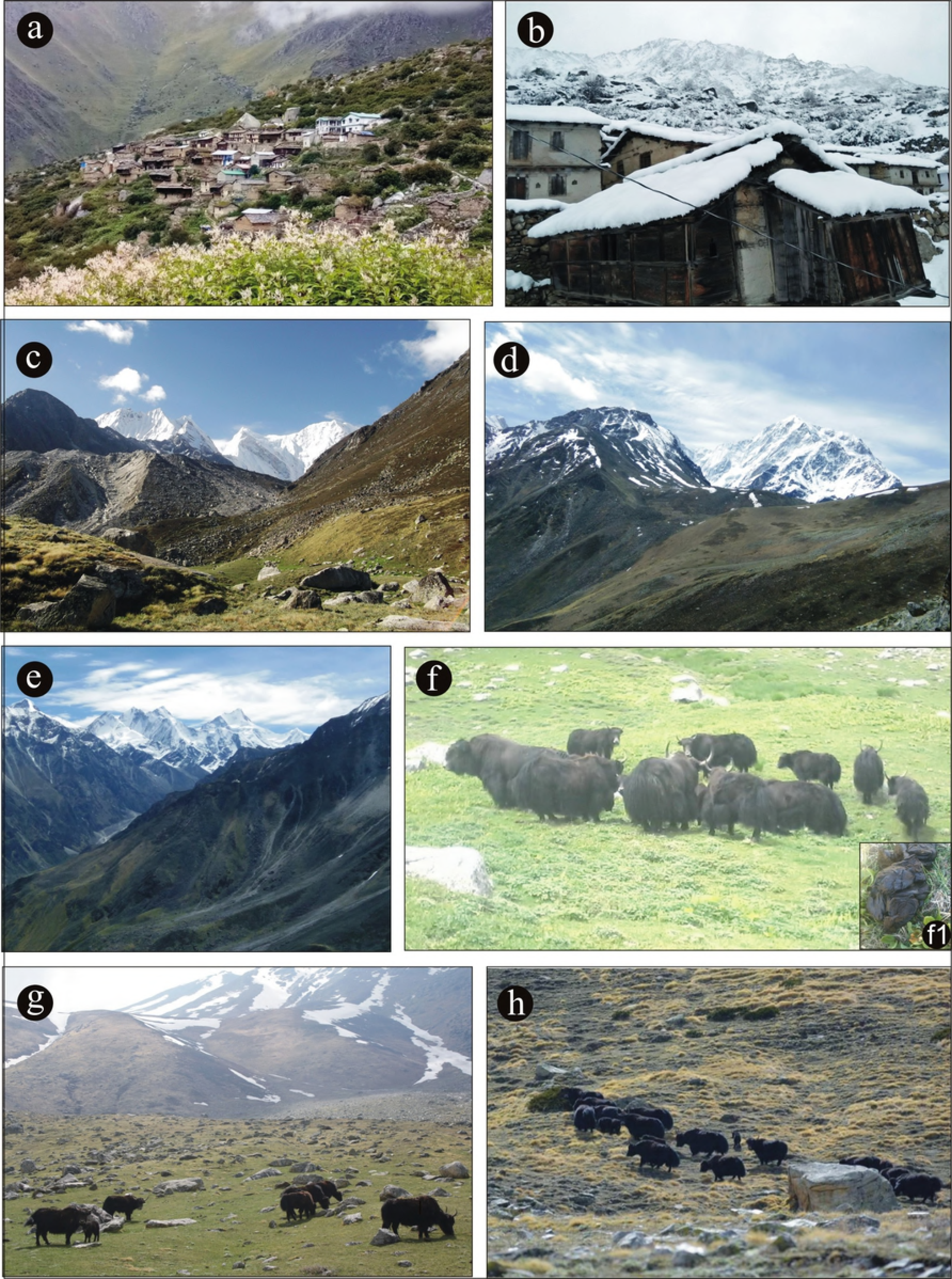
Field photographs a. Dronagiri village inhabited by Bhotiyas near Indo-Tibet border, b. A view during winter snow fall, Field photographs c. A view of Alpine meadow vegetation, d, & e. A view of Alpine steppe vegetation. Field photographs f. A group of wild Yak during resting time in Alpine meadow vegetation, g. A group of yak during grazing in Alpine steppe vegetation surrounding glacier point. Field photograph f1. Close up of Yak dung. Field photographs, h. Yak migration toward higher altitude during summer

Vegetation in the study area consists of Alpine meadow at lower elevations at 3300 meters and Alpine steppe at higher altitudes above 4000 meters (Fig 1. c., d, & e). Alpine meadow vegetation is mainly composed members of the Poaceae, Cyperaceae, and Asteraceae. Other associated taxa include members of the Polygonaceae, Onagraceae, Liliaceae, Rosaceae, Balsaminaceae, Chenopodiaceae, Amaranthaceae, and Ranunculaceae which grow luxuriantly in the region along with scattered shrubby elements, mainly *Rhododendron campanulatum, Prunus armeniaca, Juniperus squamata, Juniperus indica*, and *Rosa macrophylla*. Ferns and their allies including *Equisetum diffusum, Pteridium revolutum, Adiantum venustrum, Asplenium fortanum*, and *Lycopodium selago* are also common.

Plants characteristic of the Alpine steppe include Poaceae, *Artemisia, Carex melanantha, Caragana, Stipa orientalis*, and *Lonicera* along with scattered shrubby elements namely, *Ephedra gerardiana, Haloxylon thomsonii*, and *Capparis spinosa* [27]. The Yak is one of the mega herbivores found in the higher Himalayan region of southern central Asia, the Tibet Plateau and north Russia and Mongolia; it is considered to be critically endangered (Fig 1. f. & g.). Its preferred habitat is at high altitude, with a cool climate and generally it will tolerate temperatures as low as −40°C [28]. The climate of the region is very cold during summer and dry during winter. The maximum temperature ranges up to 12°C in summer and down to −20°C during winter. Other associated herbivorous mammals in the region include *Hemitragus jemlahicus* (Himalayan tahr) and *Moschus leucogaster* (White-bellied musk deer).

## Materials and methods

### Field work

In 2017, during the summer (March-July), the second author surveyed the site and collected11 fresh/semi-dry Yak dung samples based on their size (Fig 1. f1), each consisting of approximately 200g, from the different locations of the studied sites. Similarly, during winter (November-January), another 11 dung samples of similar size were also collected from the same areas. After the dung samples collected, they were packed separately in polythene bags to avoid contamination before laboratory processing.

### Laboratory work

The dung samples were processed for pollen using the standard acetolysis method [29]. Samples were successively treated with 10% aqueous potassium hydroxide (KOH) solution to deflocculate from the sediments, 40% hydrofluoric acid (HF) to dissolve silica, and acetolysis (9:1 anhydrous acetic anhydrite to concentrated sulphuric acid, (H_2_SO_4_) for the removal of cellulose. After that, the samples were treated twice with glacial acetic acid (GAA) and washed 3 or 4 times with distilled water. The samples were then transferred to a 50% glycerol solution with a few drops of phenol to protect against microbial decomposition. Excluding the fungal spores, 421 to 470 pollen and fern spores were counted from each sample to produce the pollen spectra. The recovered pollen taxa were categorized as arboreal taxa (tree and shrub), non-arboreal taxa (marshy and terrestrial herb), and ferns. For the identification of pollen grains, we consulted the reference slides in the Birbal Sahni Institute of Palaeosciences (BSIP) herbarium of Lucknow (India) as well as published papers and photographs [30,31].

For the diatom analysis, the samples were treated with concentrated hydrochloric acid (HCl) to dissolve carbonates and then treated with a mixture of hot nitric acid (HNO_3_) and potassium dichromate to dissolve organic materials. The samples were washed with distilled water 2 to 4 times and permanently mounted on a slide with Canada balsam for microscopic observation. The number of diatoms in the summer samples was very low and not suitable to make a proper diatom spectrum. No diatoms were observed in the winter samples. The phytoliths were observed on the same diatom slide because of the availability and clarity in the assemblage. Observation and microphotographs were done using an Olympus BX-61 microscope with DP-25 digital camera under 40X magnification. The identification of the phytoliths was based on the published literature [32].

### Statistical analysis

The statistical significance of the quantified data of pollen frequency obtained from the dung samples was determined by SPSS 11.5.0, USA. A probability of *p*-value ≤ 0.05 was taken to indicate statistical significance. The resulting data were imported into Unscrambler X Software package (Version 10.0.1, CAMO, USA) for multivariate unsupervised PCA.

### Macrobotanical analysis

For the macrobotanical analysis, 50 g of each dung sample both from summer and winter seasons were gently boiled in 200 ml of distilled water to which 5% KOH was added. After boiling, the material was sieved through a 150 μm mesh. The material was washed 2 to 4 times in distilled water and observed systematically under Stereobinocular (Leica Z6APO) microscope, and photographs were taken with a Leica DFC295 camera. Identifications were made through the consultation of published literature and reference collection of seeds and vegetative plant remains.

### FESEM-EDS analysis

The Field Emission Scanning Electron Microscope (FESEM) with Energy Dispersive Spectroscopy (EDS) analysis was also performed using FESEM (JEOL, JSM-7610F) equipped with EDS (EDAX, USA instrument) operated at 25 keV to determine the elemental composition of the yak dung.

## Results

### Microbotanical assemblage from summer dung

The 11 dung samples (S1-S11) collected from the study area were characterized by the dominance of non-arboreals (71.2%), over arboreals (25.8%). The ferns, both monolete and trilete, comprised 3.0%, of the sample. Among non-arboreal taxa, Poaceae was dominant (30.6%), followed by Cyperaceae (6.3%), and *Artemisia* (4.4%). Among arboreals the local taxa, *Rhododendron* and *Juniperus* were 3.5% and 0.9% respectively and the other extra-local arboreal taxa, *Pinus* and *Betula* were 5.0% and 2.6% (Figs 3 & 4).

Trace amounts of the diatom, *Hantzschia*, was present in all the samples. A variety of phytoliths were also observed in the same samples. Dumbbell bilobate morphology was dominant followed by elongated smooth long cell and bilobate. The others such as cuneiform, bulliform cell, rondel, and polylobate were also present in the assemblage (Fig 5).

**Fig 6.**
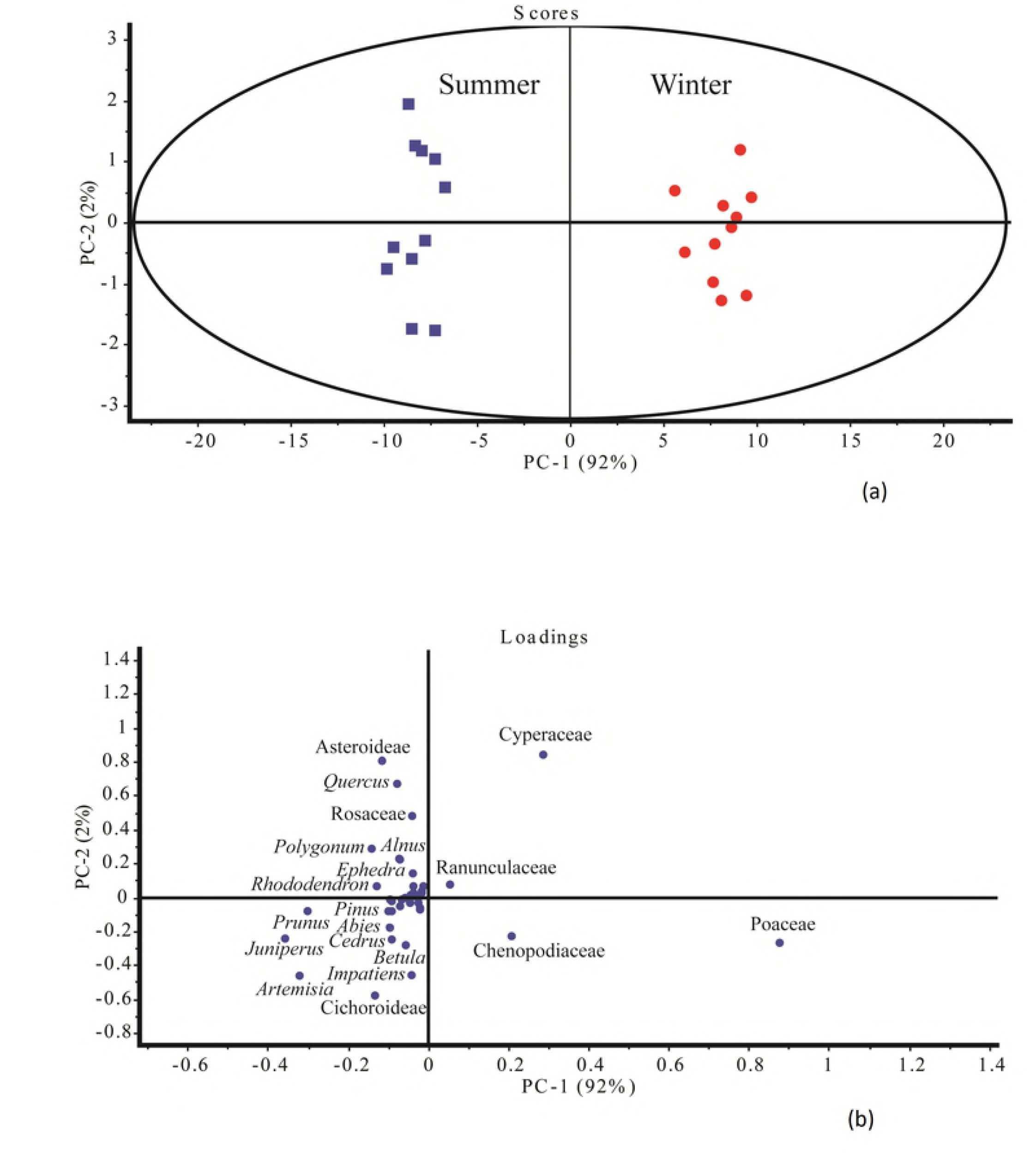
The PCA analysis of the Yak dung samples between summer (a) and winter (b) samples

**Fig 3.**
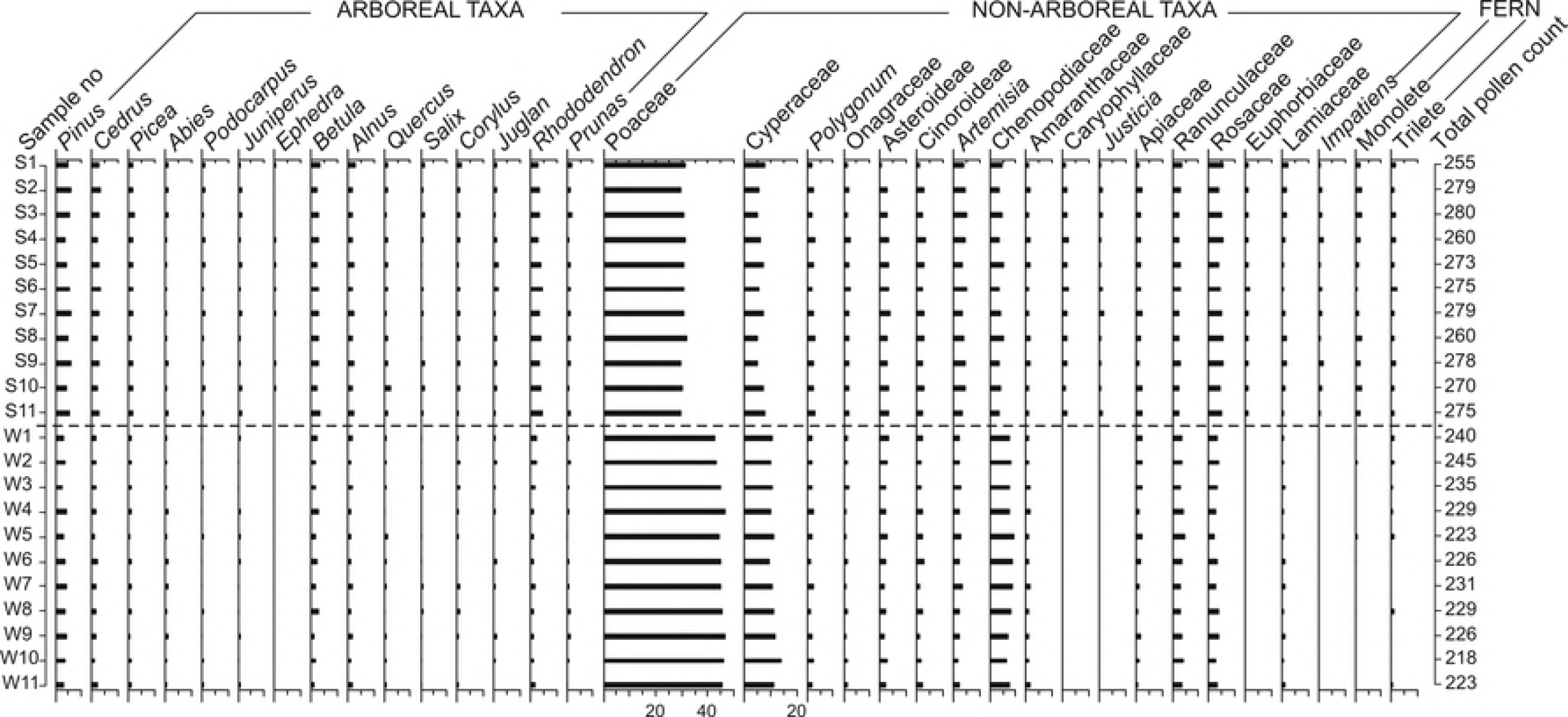
Pollen spectra of the studied Yak dung samples

**Fig 4.**
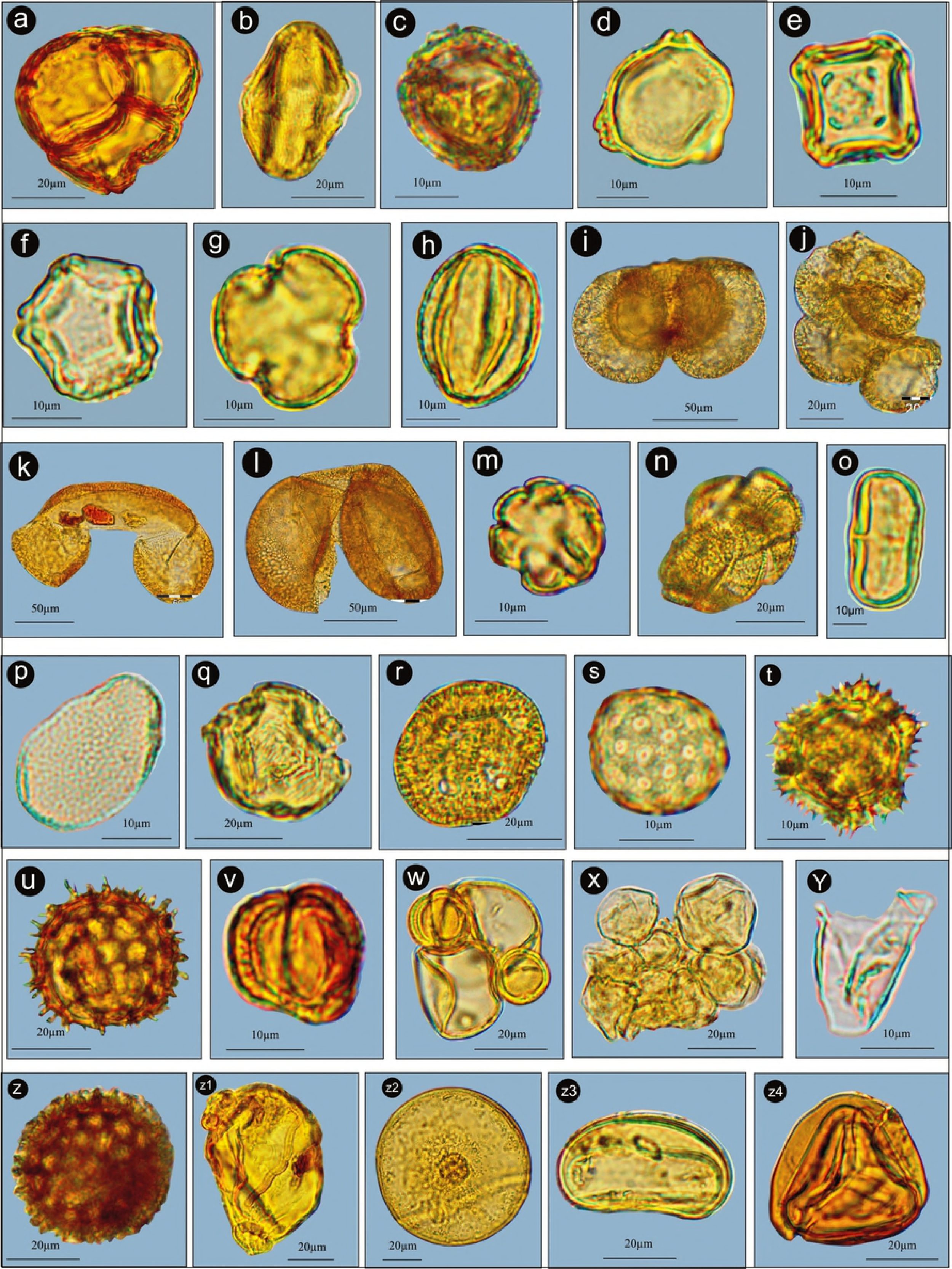
Palynoassemblage recovered from the Yak dung samples a. *Rhododendron*, b. *Prunus*, c. *Juniperus*, d. *Betula*, e. *Alnus*, f. *Corylus*, g. *Salix*, h. *Quercus*, i. *Pinus*, j. *Pinus* pollen in clumping, k. *Abies*, l. *Cedrus*, m. Euphorbiaceae, n. Convolvulaceae, o. Apiaceae, p. *Impatiens*, q. Rosaceae, r. Amaranthaceae, s. Chenopodiaceae, t. Asteroideae, u. Cinoroideae, v. *Artemisia*, w. *Artemisia* pollen associated with Poaceae, x. Poaceae pollen in clumping, y. Cyperaceae, z. *Polygonum*, aa. Onagraceae, ab. *Arcella*, ac. Monolete, ad. Trilete

### Microbotanical assemblage from winter dung

The 11 dung samples (W1-W11) collected from the study area are characterized by the dominance of non-arboreals (83.4%), over arboreals (15.8%). The ferns, both monolete and trilete, comprised 0.8%, of the sample. Among non-arboreal taxa, the Poaceae was dominant (45.0%), followed by Cyperaceae (11.0%), and Chenopodiaceae (7.8%). Among arboreals the local taxa, *Rhododendron* and *Juniperus* were represented at the value of 1.5% and 0.7% respectively. The other extra-local arboreal taxa, *Pinus*, and *Betula* were 3.5% and 1.9% respectively (Figs 3 & 4).

Diatoms were absent in all the studied samples. Phytoliths in the same samples were present with the dumbbell bilobate morphology dominant, as in the summer sample, followed by bilobate short cell and the elongated smooth long cell respectively. Other morphologies such as elongated echinate long cell, polylobate, and rondel are also regularly present in the assemblage. (Fig 5).

### PCA results

A total of 94% variance could be explained by two major pollen groups, arboreal and herbaceous taxa (Fig 6 a). The score plot showed that these two major components were responsible for the cluster differentiation. Poaceae, Cyperaceae, Chenopodiaceae, and Asteroideae were dominant and placed a high range of the PCA quadrants. The multivariate PCA and the loading plot between PC-I vs PC-II based on the differential pollen frequencies showed different pollen types responsible for cluster separation. The loading plot showed that the pollen taxa responsible for the difference between the summer and winter dung samples were Asteroideae, *Quercus*, Rosaceae, *Polygonum, Alnus*, *Ephedra*, *Rhododendron*, *Pinus*, *Prunus, Abies*, *Cedrus*, *Betula*, *Impatiens, Artemisia*, and Cichoroideae (higher in summer dung). Whereas, taxa like Poaceae, Cyperaceae, Ranunculaceae, and Chenopodiaceae were higher in winter dung samples (Fig 6 b).

**Fig 5.**
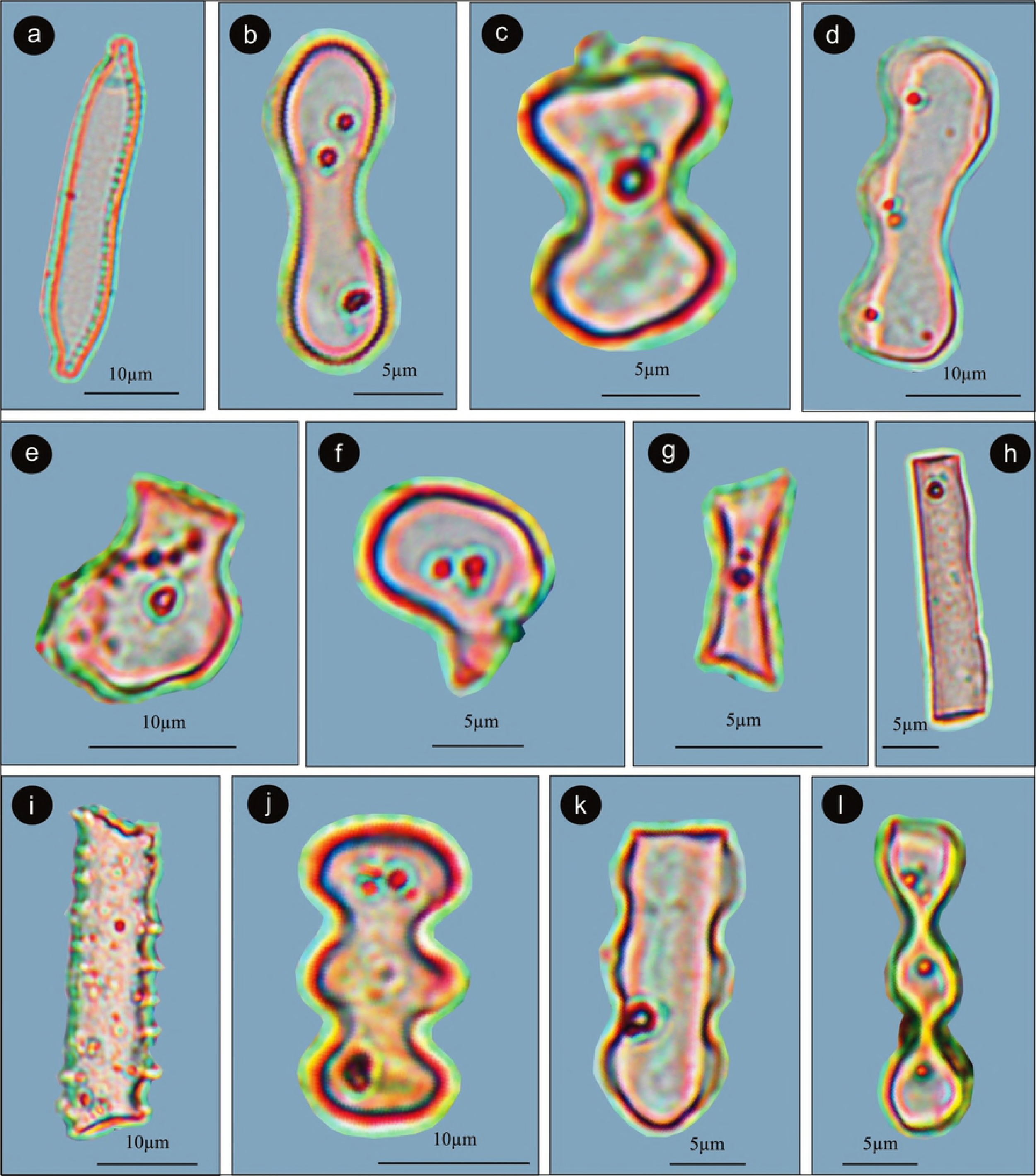
Diatom and phytoliths assemblage of the studied samples a. *Hantzschia*, b. Dumbel bilobate, c. Bilobate short cell, d. Wavy, e. Cuneiform Bulliform cell, f. Bulliform, g. Rondel, h. Elongated smooth long cell, i. Elongated echinate long cell, j. Polylobate, k. Polylobate, l. Cylindrical polylobate.

### Macrobotanical assemblage

The macrobotanical remains recovered from both summer, and winter dung samples were in good condition. Twigs and leaves of Poaceae were predominant over other herbaceous plant material from the studied samples. Monocot twigs, leaves, and dicotyledonous seeds were also present and may have become incidentally ingested along with the preferred food material. (Fig 7).

**Fig 8.**
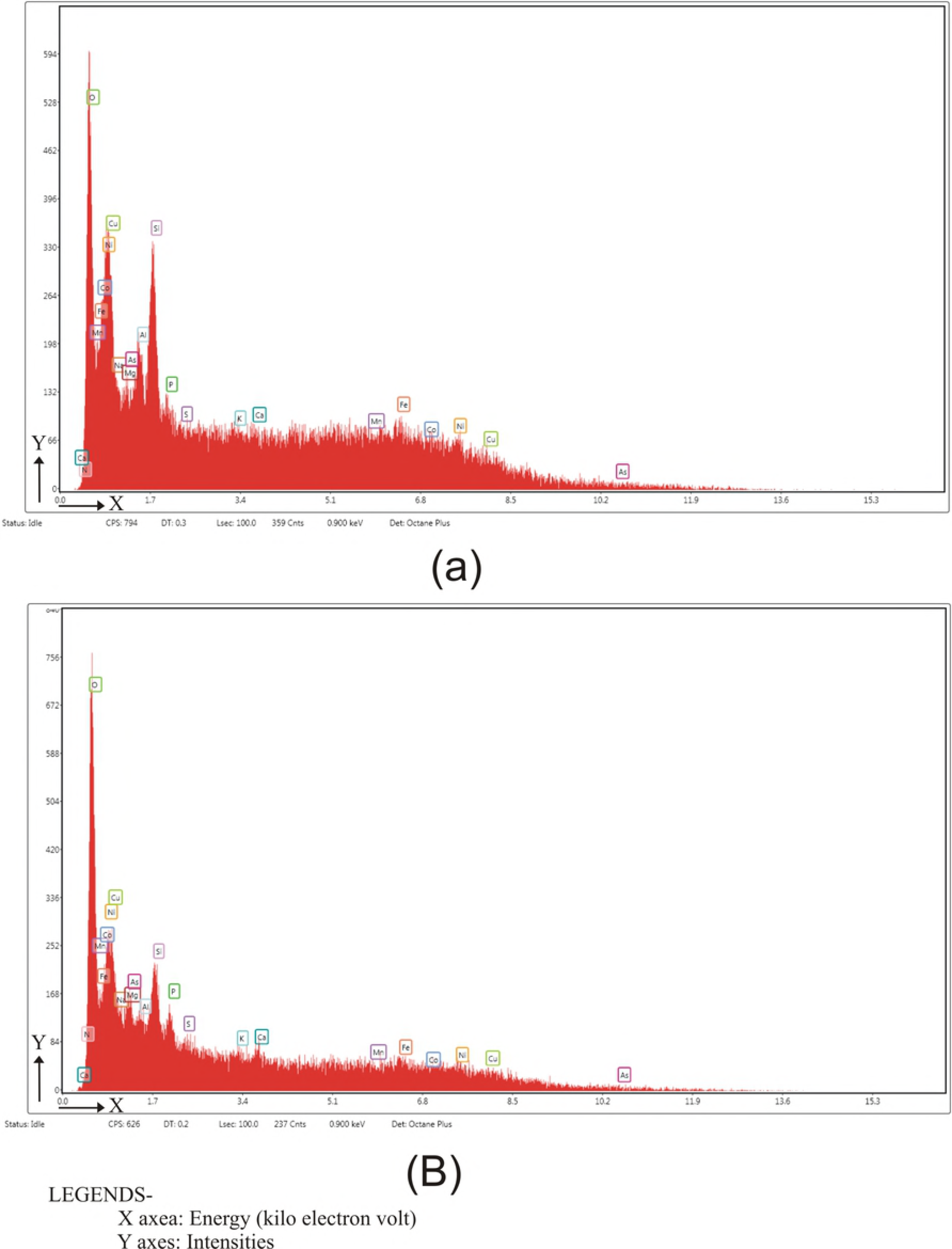
FESEM-EDS analysis micrographs in Yak dung samples collected from summer (a).

### FESEM-EDS data

The data generated from the FESEM-EDS elemental analysis of the summer yak dung samples observed that the Oxygen (O_2_) content/level is 56.89 (weight %), followed by Na, 18.95 (weight %), Si, 6.4 (weight %), Al, 4.12 (weight %), and Mg, 3.91 (weight %) Fig 8, (Table-1).

**Table 1.** List of the elements value generated by FESEM-EDS analysis in Yak dung sample collected from summer season.

| Element | Weight % | Atomic % | Error % | Net Int. | K Ratio | Z | R | A | F |
| --- | --- | --- | --- | --- | --- | --- | --- | --- | --- |
| N K | 0.04 | 0.05 | 99.99 | 0.01 | 0.0001 | 1.0631 | 0.9691 | 0.241 | 1 |
| O K | 56.89 | 68.79 | 8.79 | 64.81 | 0.2416 | 1.0436 | 0.9792 | 0.4069 | 1 |
| Na K | 18.95 | 15.95 | 14.88 | 22.41 | 0.0493 | 0.9527 | 1.0053 | 0.2726 | 1.0029 |
| Mg K | 3.91 | 3.11 | 18.28 | 6.56 | 0.0111 | 0.9703 | 1.0128 | 0.2904 | 1.0044 |
| Al K | 4.12 | 2.96 | 15.24 | 9.27 | 0.0151 | 0.9355 | 1.02 | 0.3883 | 1.006 |
| Si K | 6.4 | 4.41 | 10.22 | 18.75 | 0.0304 | 0.9571 | 1.0268 | 0.4931 | 1.006 |
| P K | 3.03 | 1.89 | 13.42 | 8.41 | 0.016 | 0.9202 | 1.0332 | 0.5685 | 1.0072 |
| S K | 1.54 | 0.93 | 12.29 | 5.08 | 0.0096 | 0.9392 | 1.0393 | 0.6605 | 1.0089 |
| K K | 1.16 | 0.57 | 19.16 | 4.01 | 0.0094 | 0.8915 | 1.0559 | 0.887 | 1.0238 |
| Ca K | 0.84 | 0.41 | 31.05 | 2.63 | 0.0073 | 0.9086 | 1.0608 | 0.9227 | 1.0283 |
| Mn K | 0.02 | 0.01 | 99.99 | 0.05 | 0.0002 | 0.8016 | 1.0816 | 1.0071 | 1.1281 |
| Fe K | 0.06 | 0.02 | 81.78 | 0.12 | 0.0006 | 0.8151 | 1.0848 | 1.0118 | 1.1614 |
| Co K | 0.03 | 0.01 | 92.07 | 0.06 | 0.0003 | 0.7973 | 1.0875 | 1.0146 | 1.2068 |
| Ni K | 0.15 | 0.05 | 68.23 | 0.25 | 0.0015 | 0.8252 | 1.0898 | 1.0162 | 1.202 |
| Cu K | 2.42 | 0.74 | 21.75 | 3.38 | 0.023 | 0.7844 | 1.0915 | 1.0168 | 1.1892 |
| As K | 0.45 | 0.12 | 67.31 | 0.31 | 0.0047 | 0.7355 | 1.0907 | 1.0133 | 1.391 |

**Fig 7.**
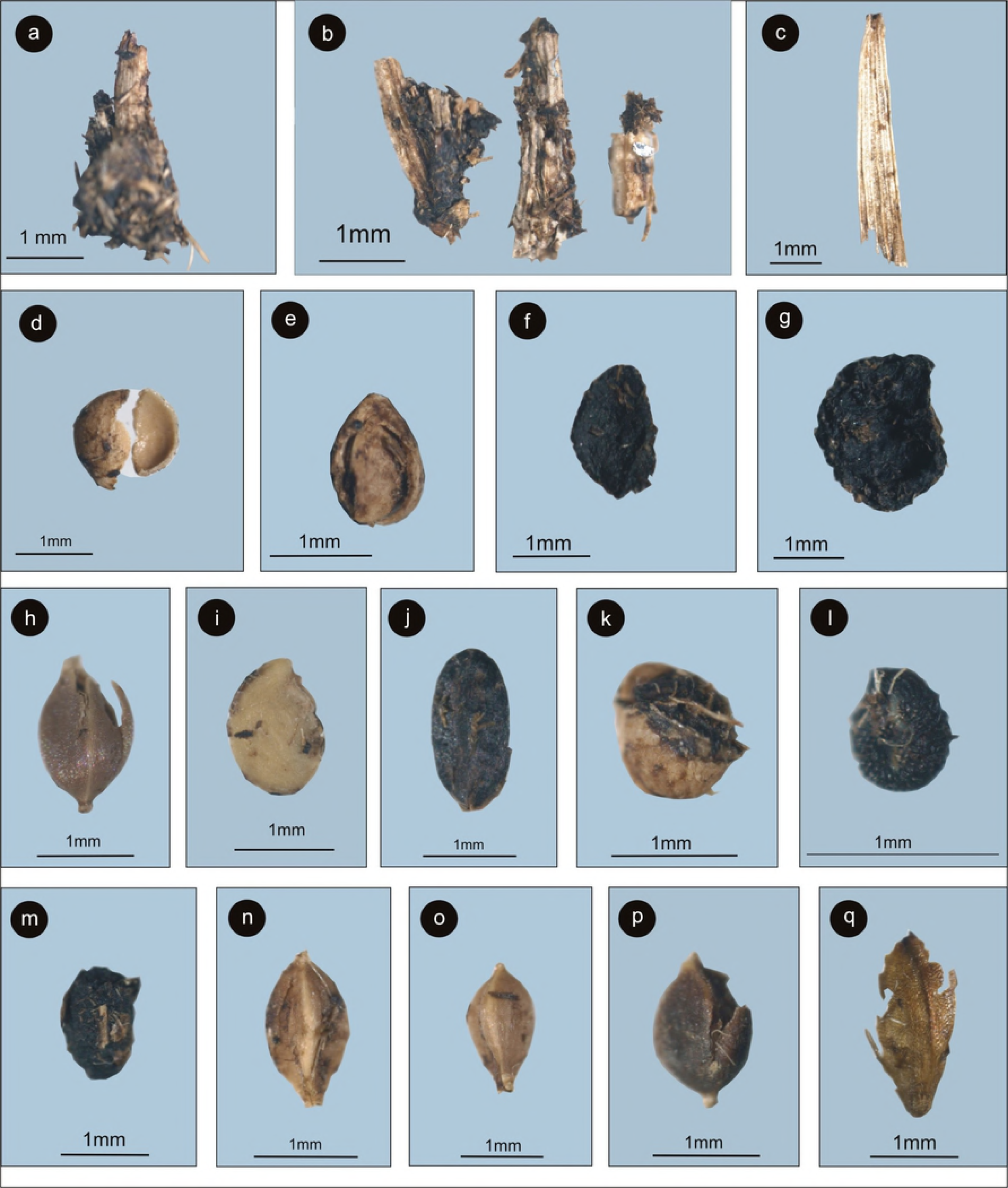
Macrobotanical assemblage recovered from the Yak dung samples a. & b. plants parts including roots and stem of sedges/herbaceous species, c. Monocot leaf, d. & e. Rosaceae, f. Liliaceae, g. *Xanthoxylum* sp., h. *Polygonum*, i. *Solanum* sp. (Solanaceae), j. Cyperaceae, k. Rosaceae, l. *Agrostema* sp., m. Fabaceae, n. & o. *Carex* sp. (Cyperaceae), p. Polygonaceae, q. Cyperaceae

The FESEM-EDS elemental analysis of the winter yak dung samples recorded that the Oxygen (O_2_) content/level is 33.82 (weight %), followed by Mn, 12.27 (weight %), Ni, 11.47 (weight %), Co, 11.12 (weight %), Fe, 10.9 (weight %), Cu, 8.76 (weight %), and Si, 8.42 (weight %) Fig. 8, (Table-2).

**Table 2.** List of the elements value generated by FESEM-EDS analysis in Yak dung sample collected from winter season.

| Element | Weight % | Atomic % | Error % | Net Int. | K Ratio | Z | R | A | F |
| --- | --- | --- | --- | --- | --- | --- | --- | --- | --- |
| N K | 0.04 | 0.07 | 99.99 | 0.01 | 0.0001 | 1.1582 | 0.9016 | 0.2556 | 1 |
| O K | 33.82 | 61.84 | 12.31 | 34.38 | 0.1513 | 1.1388 | 0.9129 | 0.3928 | 1 |
| Na K | 0.02 | 0.03 | 99.99 | 0.01 | 0 | 1.0439 | 0.9428 | 0.1259 | 1.0017 |
| Mg K | 0.01 | 0.01 | 99.99 | 0.01 | 0 | 1.0644 | 0.9518 | 0.1987 | 1.003 |
| Al K | 0.32 | 0.35 | 86.52 | 0.5 | 0.001 | 1.0273 | 0.9603 | 0.2906 | 1.0054 |
| Si K | 8.42 | 8.77 | 15.33 | 18.98 | 0.0363 | 1.0521 | 0.9684 | 0.4075 | 1.006 |
| P K | 0 | 0 | 99.99 | 0.01 | 0 | 1.0125 | 0.9762 | 0.4766 | 1.0099 |
| S K | 0 | 0 | 99.99 | 0 | 0 | 1.0343 | 0.9836 | 0.5938 | 1.0152 |
| K K | 0.19 | 0.14 | 66.28 | 0.59 | 0.0016 | 0.9841 | 1.0042 | 0.8487 | 1.0523 |
| Ca K | 0.71 | 0.52 | 56.3 | 2.1 | 0.0068 | 1.0037 | 1.0105 | 0.8968 | 1.0727 |
| Mn K | 12.27 | 6.53 | 7.82 | 25.14 | 0.125 | 0.8887 | 1.0381 | 0.9967 | 1.1507 |
| Fe K | 10.9 | 5.71 | 8.44 | 19.65 | 0.1098 | 0.9044 | 1.0427 | 0.9996 | 1.1141 |
| Co K | 11.12 | 5.52 | 10.06 | 16.57 | 0.104 | 0.8853 | 1.047 | 0.9763 | 1.0826 |
| Ni K | 11.47 | 5.72 | 9.31 | 15.27 | 0.1068 | 0.917 | 1.0508 | 0.9624 | 1.0547 |
| Cu K | 8.76 | 4.03 | 14 | 9.62 | 0.0771 | 0.8725 | 1.0542 | 0.9529 | 1.0592 |
| As K | 1.95 | 0.76 | 59.62 | 0.98 | 0.0174 | 0.8216 | 1.0612 | 0.9687 | 1.121 |

## Discussion

The micro and macrobotanical remain preserved in the summer and winter yak dung, and their elemental analysis indicates seasonal differences in the yak’s dietary preference in relation to the vegetation composition and climate in the region. The overall pollen data of both the summer and winter collected yak dung samples indicates that grasses form the primary component of their diet. The other associated herbs and shrubs, Cyperaceae, *Artemisia*, Asteroideae, Chenopodiaceae, *Impatiens, Prunus*, and *Rhododendron* are also very important parts of the species diet. It is also observed that yak prefer plant diversity in their diet which is reflected in the summer samples as demonstrated by the higher pollen diversity and Yak will migrate up to 50 kilometres to forage for their food [33, 34].

The presences of Cyperaceae, Rosaceae, Ranunculaceae, and Polygonaceae in the palynoassemblage were also identified in summer and winter seasons and are characteristic of Alpine meadow vegetation in the region. The grasses recovered are closely associated with Rosaceae, Fabaceae, Asteraceae, and Lamiaceae which dominate in Alpine meadow vegetation regarding cover and abundance. Alpine steppes are characterized by high percentages of Chenopodiaceae, and *Artemisia* along with *Ephedra* and *Nitraria* [35]. The association of *Artemisia, Caraguna, Ephedra, Juniperus, Salix*, and *Lonicera* is characteristic of the Alpine steppe vegetation in the higher portions of the Himalaya [36]. As our studied samples also included *Ephedra, Juniperus*, Chenopodiaceae, and *Artemisia* pollen in the palynoassemblage and complements the presence of Alpine steppe vegetation in and around the study region. The presence of *Ranunculus* was noticeable in the studied palynoassemblage and found to be characteristic taxa of snow bed vegetation [37]. The presence of Cyperaceae, Polygonaceae, and Onagraceae along with *Botryococcus, Hantzschia*, and *Arcella* seen in the dung samples are indicative of the perennial water-logged condition and streamlets running in and around the study areas. Their absence in the winter samples reflects the absence of unfrozen water. The recovery of shrubby taxa, *Rhododendron* and *Prunus*, along with *Impatiens* and Euphorbiaceae in the palynoassemblage was observed and are strongly suggestive of the monsoonal activity in the region [12,38]. The recovery of Chenopodiaceae and *Artemisia* is very significant as these taxa are strongly indicative of the winter dryness in the region. The abundance of *Artemisia* pollen in the winter palynoassemblage indicates the seasonal glacial condition/phase in the region [39] which is reflected in our studied samples. However, many workers observed this distinctive assemblage in surface soil samples and had utilized the Chenopodiaceae/*Artemisia* ratio to determine the relative dryness of the Alpine steppes and deserts in the region, so is a useful climatic indicator [35, 40–43] which suggest that a key role can be provided by studied dung samples. But to date, no such work has been initiated on dung samples.

Among arboreal taxa, the presence of *Pinus, Cedrus*, and *Alnus* in the palynoassemblage which do not grow in this region are indicative of upthermic wind activity which transported this pollen from the conifer forest zone present at lower elevations,1000 to 3300 meters. Pollen from these taxa in the dung samples would have been incorporated through secondary ingestion of the food plant, exposed dung, and soils in the region on which they would have settled. Likewise, the low value of *Juniperus* pollen would have also been incorporated through the ingestion of the plants. *Rhododendron* pollen is entomophilous, it presence in the assemblage would have been incorporated through the ingestion of the plant’s flowers and must have been local in origin. The presence of *Salix* and *Prunus* pollen in the palynoassemblage is indicative of the presence of perennial water channels, streams and moist condition in the region as these plants generally grow along water channels and moist places [35,37,44]. Similarly, the presence of fern spores, both monolete and trilete, in the palynoassemblage is indicative of the warm and humid condition in the region during the summer.

The diatom and phytolith analysis from both summer and winter dung samples revealed that the only a few diatom taxa were recovered in the summer dung samples. The presence of diatoms is suggestive of at least seasonal perennial water in the region which became incorporated in the dung through the ingestion of the water. Among phytoliths, grass phytoliths predominant as are the macrobotanical remain of grasses while the other phytoliths also constantly represented in the assemblage and are also indicative of plants other than grasses that were also important food plants of the yak.

The FESEM-EDS elemental analysis of Yak dung samples was also conducted to understand the elemental percentage in relation to the vegetation composition and climate in the region. Sixteen elements have been identified and characterized in the yak dung samples (Table 1 & 2). As the large animals play an important role in the nutrient cycles due to their ability to travel long distances [25,26], large-bodied forms such as the yak can perform a critical role in this cycle as indicated by the elemental values seen in the dung samples. The redistribution of these elements would play an important role for both flora and fauna in the region. For example, the distribution of phosphorous and sodium have been identified as having an important role in the extinction of the both flora and fauna in the region [15]. These elements in the yak dung samples could be useful to understand the nutritional value in relation to the vegetation composition and the species dietary preferences in the region.

The macrobotanical assemblage (Cyperaceae, Polygonaceae, and Rosaceae) was in good agreement with pollen data in dung samples in relation to the current vegetation, and climate in the region (Fig 7).

### Biodiversity of summer and winter dung samples

There are some differences between the summer and winter dung samples in the palynoassemblages. The pollen diversity in summer samples is comparatively higher than the winter samples. In summer, the forage is relatively abundant and nutritious and the yak move up to higher altitudes and occupy a wider area including both Alpine meadow and steppe vegetation regions (Fig 1, h). During the winter season, they may either moves towards lower elevations or remain in the higher mountain sides with minimal movement reflecting the scarcity of forage due to snow cover at high altitude. Most of the plant taxa, especially arboreal blooms in the summer season and therefore the diversity of pollen taxa in summer dung samples are always higher than winter dung samples. The presence of *Botryococcus, Arcella*, and *Hantzschia* in the summer samples was higher than in the winter samples reflecting the greater availability of free-flowing water. In the elemental analysis, the silica content is comparatively higher in the winter samples in response to the dominance of grass pollen.

It is observed that the elemental contents of summer samples were different than the winter samples. The Na, Mg, and Al content were low in the winter samples, and the reason may be due to the scarcity of food plants. Phosphorous and sulfur are absent in the winter samples. However, the O, Mn, Ni, Co, Cu, and Si content in the winter sample were relatively higher than the summer sample. The diversity of the seeds, fruits and twigs were comparatively more common in the winter samples as the maximum occurrences of flowering, and fruiting and the twig of the shrubby elements are more due to being consumed by the yak during that time.

### Statistical significance of pollen frequencies

A critical review of the PCA indicates a clear seasonal periodicity in the appearance of the pollen of different plant species especially the herbaceous taxa, which were in full bloom during the winter season. A clear-cut seasonal clustering of these pollen taxa is present and mainly represented by two major pollen groups, arboreals and herbs. The major arboreal group like *Pinus, Abies, Cedrus, Betula, Quercus, Rhododendron*, and *Prunus* were recorded in the summer samples of yak dung, attributed to their maximum blooming in spring season (February to March) and further deposition on herbaceous vegetation, and could be secondarily incorporated as yak’s diet. The dominance of Poaceae, Ranunculaceae, and Chenopodiaceae in winter dung could coincide with their peak flowering during the winter season. It was also clear from the loading plot that maximum diversity of plant taxa in yak dung was in summer season owing to their activities in summer where they cover a larger range. However, in winter during snow fall, they tend to be more inactive finding place for their sustenance, with a dependence on primary herbaceous plant taxa (Fig 6).

### Linkages with endangered and extinct megaherbivores

A study conducted on stomach contents including pollen and spores from fossil woolly rhinoceros (*Coelodonta antiquitatis*) from Russia [13] revealed the presence of predominately non-arboreal pollen taxa including Poaceae and other associated herbs (98.5%), followed by arboreal pollen (trees and shrubs) (0.9%) and spores (including ferns and mosses) (0.6%) respectively. Our dataset for the pollen assemblage from the yak dung samples is closely similar to that of the woolly rhino stomach contents pollen fossil data. Similarly, a study conducted on the mammoth diet based on dung indicates a similarity between these two megaherbivorous mammals, with Poaceae in both taxa as the primary component of their diet followed by the Cyperaceae and Asteraceae. The absence or only trace amounts of arboreal pollen taxa is indicative of the grassland vegetation utilized by the mammoth which lived in cold climatic conditions at higher latitudes [37], that were comparable to the vegetation of the higher elevations at which the yak lives today.

It should be noted that during the Pleistocene fossil yak has been found in eastern Russia, Tibet, and Nepal [45] and so was directly associated with woolly mammoths, and most likely also the woolly rhino. The woolly rhinoceros and mammoth became extinct in Eurasia because of landscape changes during Pleistocene-Holocene boundary (12000-9000 years BP) due to the formation of widespread forest in the temperate and arctic regions of northern Eurasia and loss of grassland [46]. Yak and Bison share a common ancestry and both originated in central Asia, and mitochondrial DNA analysis also suggests that the yak is closely similar to Bison [47,48], but they are distinct geographically. While the yak remained restricted to western Asia, bison dispersed westward into Europe and northeast across the Bering Land Bridge into North America during the middle to late Pleistocene [45, 49, 50–52]. Fossil yak has been reported from Alaska, but radiocarbon dates of these specimens have shown that they are of domestic cattle brought in by miners [53]. The extinct bison (*Bison priscus*) is associated with yak in Eurasian faunas. Studies of its diet based on microhistological fecal analysis indicate that like the mammoth and woolly rhinoceros 98% of its diet was grasses, followed by Cyperaceae [54]. However, unlike these faunas, the diet of bison changed between the late Pleistocene to early Holocene which may have permitted the survival of the recent bison (*Bison bison*) in North America and the wisent (*Bison bonasus*) in Europe [55].

This data might be helpful in understanding the extinction of megaherbivorous animals such as woolly rhino and mammoth despite having diets very similar to the surviving yak and bison based on their similar palynodata. There are differences between the woolly rhino and today’s living one horn rhino regarding habit and diet. A study of the pollen and non-pollen palynomorphs preservation in the dung of the extant one horn rhino [12, 56] reveals that while grass is the primary diet of both species, the living rhino also required perennial water-logged conditions and a flood plain area that included both large grasslands along with scattered woodland.

## Conclusions

The multiproxy data presented here indicate that the yak utilizes a combination of both Alpine meadow and steppe vegetation depending on the season. Its response to the seasonally cold climate is either by moving to lower elevations or minimizing movements at higher elevations which have a more limited food supply. A critical part of its habitat appears to be the presence of a perennial water system. So, this documented data might serve as a strong proxy to interpret vegetation and climatic shifts in the higher Himalaya and to correlate them at a global level.

Climate change with warmer average temperatures may extend the length of time that unfrozen water may be available throughout the region, providing the yak with an opportunity to extend its range. However, as a species adapted to cold temperatures, any increase in mean summer temperatures may force the species to spend more time at higher elevations, above the lusher Alpine meadows, thus reducing access to a major source of food, and also reducing its overall range.

The dung of herbivorous mammals can provide a durable substrate that allows the investigation of the modern pollen data that complements the data recovered from modern surface soil in relation to the vegetation composition and climate in the region. The study of macrofossils, diatoms, and phytoliths can be used to supplement the pollen database to prevent incorrect interpretation for the palaeodietary analysis, when wind transported pollen such as *Pinus, Cedrus*, and *Abies* are present in the assemblage. The elemental analysis of different elements in the dung sample also provides a better understanding of the relationship between the surface soil samples and vegetation composition in the region. This multiproxy dataset can help to understand the collapse of species and the subsequent extinction of the megafauna from the different region of the world. The generated data will be helpful for the differentiation of the temperate and tropical megaherbivorous animals in relation to the database. The diet of wild Yak existing in the western Himalayas also includes the consistent occurrence of some secondary herbs and trees beside, grasses as primary food, thus this flexibility in dietary habit could be one of the reason for their survival through Pleistocene-Holocene vegetation transition where other megafaunas become extinct.

While it is clear that, yak, bison, rhinos and mammoth, were capable of feeding on the same types of vegetation, predominately grasses, and so had similar preferences in their diet, there were significant differences in their preferred habitat. The wild yak survives today, although with a much-reduced distribution. Its current distribution corresponds to the combination of the vegetation composition and colder climate that exists at the higher elevations in the Himalayan region. Possibly due to their larger size and need for greater quantities of food the larger mammoth and rhino were not able to make the transition to, this habitat. The closest living relative to the woolly rhino is *Dicerorhinus sumatrensis*, with the genus *Rhinoceros (unicornis* and *sondaicus*) forming their sister group [57]. This raises an interesting paradox since the high elevations of the Tibetan Plateau has been proposed as the area of origin of the woolly rhino [58]. In contrast the bison survived by a change in its diet. So, while this diversity of grazers all lived at the same time and shared a common habitat, they reflect the three ways to respond to climate change; track the preferred habitat, adapt to changing conditions or go extinct.

## Acknowledgements

We thank, Director, Birbal Sahni Institute of Palaeosciences, Lucknow, India for encouragement and permission to publish the paper (No. BSIP/RDCC/Publication no. 19/2018-19). We are thankful to Dr. Subodh Kumar for FESEM-EDS studies. We are also thankful to Mr. Amar Singh, Department of Veterinary, Chamoli District, Uttrakhand for his help during samples collection. We are thankful to Mr. Jagdish Prasad for sample maceration and Mrs. Tusha Tripathi for her kind help.

## Author contributions

S.K.B. conceived the ideas, analysed the data, and led the writing; H.S., conceived the ideas, collected samples and writing; H.G.M., analysed the data and led the writing; S.T. analysed the data, and led the writing; A.K.P. analysed the data, and led the writing.

